# A chemical-genetic system to rapidly inhibit the PP2A-B56 phosphatase

**DOI:** 10.1101/2024.07.23.604754

**Authors:** LA Allan, A Corno, JM Valverde, R Toth, T Ly, AT Saurin

**Author notes:** Equal contribution (LA Allan, A Corno, JM Valverde).

## Abstract

Serine-threonine phosphatases have been challenging to study because of the lack of specific inhibitors. Their catalytic domains are druggable, but these are shared or very similar between individual phosphatase complexes, precluding their specific inhibition. Instead, phosphatase complexes achieve specificity by interacting with short-linear motifs (SLiMs) in substrates or their binding partners. We develop here a chemical-genetic system to rapidly inhibit these interactions within the PP2A-B56 family. Drug-inducible recruitment of ectopic SLiMs (“directSLiMs”) is used to rapidly block the SLiM-binding pocket on the B56 regulatory subunit, thereby displacing endogenous interactors and inhibiting PP2A-B56 activity within seconds. We use this system to characterise PP2A-B56 substrates during mitosis and to identify a novel role for PP2A-B56 in maintaining kinetochore-microtubule attachments at metaphase. The directSLiMs approach can be used to inhibit any other phosphatase, enzyme or protein that uses a critical SLiM-binding interface, providing a powerful strategy to inhibit and characterise proteins once considered .“undruggable”.

## Introduction

The PPP family of serine-threonine phosphatases dephosphorylate the majority of phospho-Ser/Thr residues in cells (1, 2). This large family of enzymes have been notoriously difficult to study because they share the same or very similar catalytic subunits, precluding their specific targeting by small molecule inhibitors (3). Natural toxins have been identified that display some preference for either the PP1 or PP2A catalytic domains (4), but these domains are still shared between multiple different holoenzyme complexes. For example, PP2A exists as a heterotrimeric phosphatase containing a PP2A catalytic domain, a scaffold domain, and one of four different classes of divergent regulatory subunits - B’ (B55, PR55), B’’ (B56, PR61), B’’’ (PR48/PR70/PR130), and B’’’’ (PR93/PR110 or Striatins) (5). It is protein-protein interactions mediated by these regulatory subunits that target PP2A to specific substrates. Each regulatory subunit is present as multiple different isoform variants, further increasing diversity within the PP2A family. PP2A-B56 holoenzymes, for example, use six different B56 isoforms encoded by separate genes (6).

A major substrate binding pocket on B56, which is well conserved in all B56 isoforms, binds to short-linear motifs (SLiMs) containing a consensus LxxIxE sequence (7). A wide variety of PP2A-B56 substrates contain LxxIxE motifs, and their presence within conserved unstructured regions of proteins can be used to predict PP2A-B56 substrates (7, 8). LxxIxE motifs often contain a serine or threonine residue within or immediately after this motif, and phosphorylation of these residues can enhance B56 binding strength (9-12). This allows phosphorylation inputs to recruit PP2A-B56 to specific substrates or locations at certain times. For example, CDK1 and PLK1-dependent binding of PP2A-B56 to BUBR1 allows this interaction to occur specifically at kinetochores during mitosis, where it is needed to promote dephosphorylation of sites that would otherwise impede microtubule binding (13, 14). This is important to stabilise kinetochore-microtubule attachments and allow proper chromosome segregation.

Mutation of specific LxxIxE motifs within substrates has been crucial for characterising PP2A-B56 function. For example, mutation of the LxxIxE motif in BUBR1 uncovered roles for PP2A-B56 in promoting kinetochore-microtubule attachments and antagonising the spindle assembly checkpoint (9, 10, 15-22). Similar strategies have been used to identify roles for PP2A-B56 in cytokinesis via RacGAP1 binding (7), and homologous recombination via BRCA2 binding (23).

A complementary approach to characterise PP2A-B56 binding partners and substrates is to mutate the LxxIxE-binding pocket on B56 (hereafter referred to as the substrate binding pocket). Mutations of key residues in B56 can prevent LxxIxE motif binding, and this was used to identify a role for this binding pocket in promoting Sgo1-binding to protect cohesion during mitosis (24). This type of approach can also be used to assess PP2A-B56 function more globally, for example to look for binding partners and phosphorylation events that change when the substrate pocket is mutated. However, it takes days to switch to a mutant form of B56, during which time cells accumulate in mitosis with unattached kinetochores due to lack of BUBR1 binding. This complicates the assessment of PP2A-B56 function during later mitosis or at any other cell cycle stage.

An alternative approach was developed by the Nilsson lab to block the substrate binding pocket by the inducible expression of a high-affinity tetrameric LxxIxE peptide (25). This was shown to bind and occlude the substrate binding pocket on B56, allowing the identification of novel substrates during mitosis or S-phase (25). However, it still takes 16 hours to express such a protein to high enough levels to inhibit B56 function, complicating interpretations about whether the resulting phospho-proteomic changes are direct or indirect. A method to more rapidly inhibit the substrate binding pocket on B56 would allow acute inhibition of enzyme activity during any cell cycle stage or any process, thus facilitating the analysis of PP2A-B56 function.

Here we develop a chemical-genetic system to inducibly recruit an LxxIxE-containing peptide to block the B56 substrate-binding pocket within seconds of small molecule addition. This can rapidly compete off B56 substrates and inhibit PP2A-B56 function, as evidenced by the fact that increased substrate phosphorylation is observed in as little as 15 seconds. We use this to comprehensively characterise acute phosphorylation changes proteome-wide following PP2A-B56 inhibition during mitosis, and to identify new roles for PP2A-B56 in maintaining stable kinetochore-microtubule attachment at metaphase. We name this approach directSLiMs - for **d**rug-**i**nducible **r**ecruitment of **ect**opic **SLiMs** - because it is a simple yet powerful approach to block critical SLiM-based interactions on other phosphatases (3), enzymes (26) or even non-enzymatic proteins (26, 27).

## Results

We set out to develop a rapid drug-inducible recruitment strategy to block the LxxIxE binding pocket on B56 and thereby inhibit substrate dephosphorylation (Figure 1A). We chose to use the FKBP-FRB interaction that can be induced by rapamycin or rapamycin analogues (rapalogs) (28), to recruit a non-phospho-dependent LxxIxE peptide, initially characterised from Ebola (**L**PT**I**H**E**EEEE: hereafter named LIE1), to the PP2A-R1A scaffolding subunit (8). We used a ribosome-skipping T2A sequence to produce separate LIE1^FRB^ and ^FKBP^R1A proteins from the same doxycycline-inducible plasmid that was integrated into a FRT site within HeLa-FRT cells (Figure 1B). Figure 1C shows that this plasmid can be used to replace endogenous R1A subunit with comparable levels of ^FKBP^R1A. Anti-FLAG immunoprecipitations shows that the flag-tagged ^FKBP^R1A subunit is able to bind to the PP2A-catalytic domain and various B56 regulatory subunits, indicating association into PP2A-B56 holoenzyme complexes. Rapamycin treatment for 30 mins, to induce recruitment of the co-expressed LIE1^FRB^, did not perturb PP2A-B56 holoenzyme assembly, but it did efficiently compete off B56-binding partners that are known to interact via the LxxIxE binding pocket (Figure 1C: GEF-H1, BubR1, RepoMan) (7).

PP2A-B56 is recruited to kinetochores during mitosis by binding to an LxxIxE motif in the kinetochore attachment regulatory domain (KARD) of BUBR1 (amino acids 664-681) (9, 10). From this position it regulates many kinetochore phosphorylation events, including the adjacent BUBR1-pT620, which is a phospho-dependent recruitment site for Polo-like Kinase 1 (PLK1) (Figure 1D) (17, 18, 29, 30). A 20-min treatment with rapamycin was sufficient to displace ^FKBP^R1A from the kinetochore and to increase BUBR1-T620 phosphorylation in LIE1^FRB^ cells (Figure 1E). This required the LIE1 peptide to bind the substrate binding pocket on B56, because rapamycin had no effect in ^FKBP^R1A cells expressing an AAA^FRB^ mutant that is analogous to LIE1^FRB^ but contains alanine residues in key B56 binding positions (Figure 1E). In summary, drug-inducible recruitment of the ectopic SLiM LIE1 (directSLiM^LIE1^), to the R1A subunit competes off B56 substrates and displaces PP2A-B56 from the kinetochore, leading to increased phosphorylation of the well-established kinetochore substrate BUBR1.

**Figure 1.**
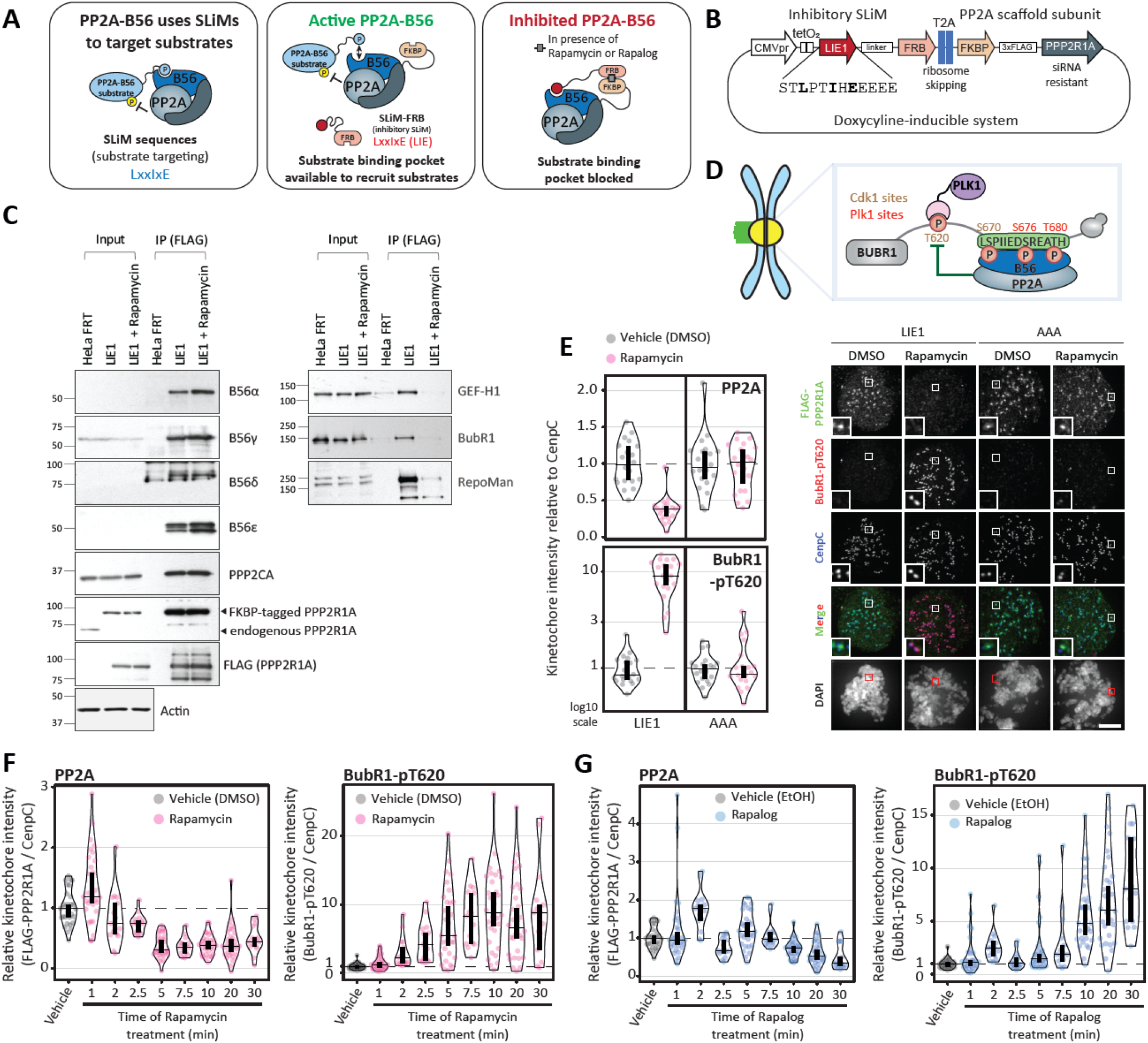
A chemical-genetic system to rapidly inhibit PP2A-B56. **A**. Schematic illustrating the principle of the drug-inducible recruitment of ectopic SLiMs (directSLiMs) system. **B**. Representation of the plasmid used to implement the directSLiMs strategy in HeLa FRT cells. **C**. Immunoblot of the PP2A complex (left column) and PP2A substrates (right column), following FLAG immunoprecipitation from nocodazole-arrested HeLa FRT cells with/without directSLiM^LIE1^ expression, -/+ rapamycin for 30min. Representative of two experiments. **D**. Schematic illustrating how PP2A-B56 regulates the phosphorylation of the Thr620 site on BUBR1. **E**. Effects of PP2A-B56 inhibition on the levels of FLAG-PPP2R1A and BUBR1-pT620 at unattached kinetochores in nocodazole-arrested HeLa FRT cells expressing the directSLiMs^LIE1/AAA^ and treated with vehicle or rapamycin for 20min. Left panel: kinetochore intensities from 20 cells, 2 experiments. Right panel: representative example immunofluorescence images of the kinetochore quantifications shown in the left panel. The insets show magnifications of the outlined regions. Scale bars: 5μm. Inset size: 1.5μm **F-G**. Effects of PP2A-B56 inhibition on the levels of FLAG-PPP2R1A and BUBR1-pT620 at unattached kinetochores, in nocodazole-arrested HeLa FRT cells expressing directSLiM^LIE1^ and treated with rapamycin (F) or rapalog (G) for 0-30 mins. Kinetochore intensities from 20 cells, 2 experiments. Data information: Kinetochore intensities in E-G are normalized to LIE1 vehicle condition. Violin plots show the distributions of kinetochore intensities between cells. For each violin plot, each dot represents an individual cell, the horizontal line represents the median and the vertical one the 95% CI of the median, which can be used for statistical comparison of different conditions (see Methods).

To examine how quickly directSLiM^LIE1^ could inhibit PP2A-B56 we performed similar analysis after 0-30 mins of rapamycin treatment. This demonstrated that PP2A delocalisation and enhanced BUBR1 phosphorylation occurred within 1-2 minutes of rapamycin addition (Figure 1F). We also used a rapamycin analogue (also known as A/C heterodimeriser, but hereafter called rapalog) that binds to the FRB-T2098L mutant present in the LIE1^FRB^, but not to endogenous FRB, thus preventing inhibition of the endogenous TORC1 complex (31). Rapalog treatment similarly inhibited PP2A-B56, although the timescale of inhibition was slightly delayed (by 5-10 mins), perhaps due to differences in FRB-FKBP affinity and/or rapalog cell permeability (Figure 1G).

To globally assess phosphorylation changes following PP2A-B56 inhibition, nocodazole-arrested mitotic directSLiM^LIE1^ cells were treated for 30 min with DMSO, rapamycin or rapalog, followed by TMT labelling, phosphopeptide enrichment and measurement by mass spectrometry (MS) (Figure 2A). A total of 9,907 phospho-sites were quantified and of these 187 exhibited a significant >1.5-fold increase following rapamycin or rapalog treatment (Figure 2B, Supplementary Figure 1A).

**Figure 2.**
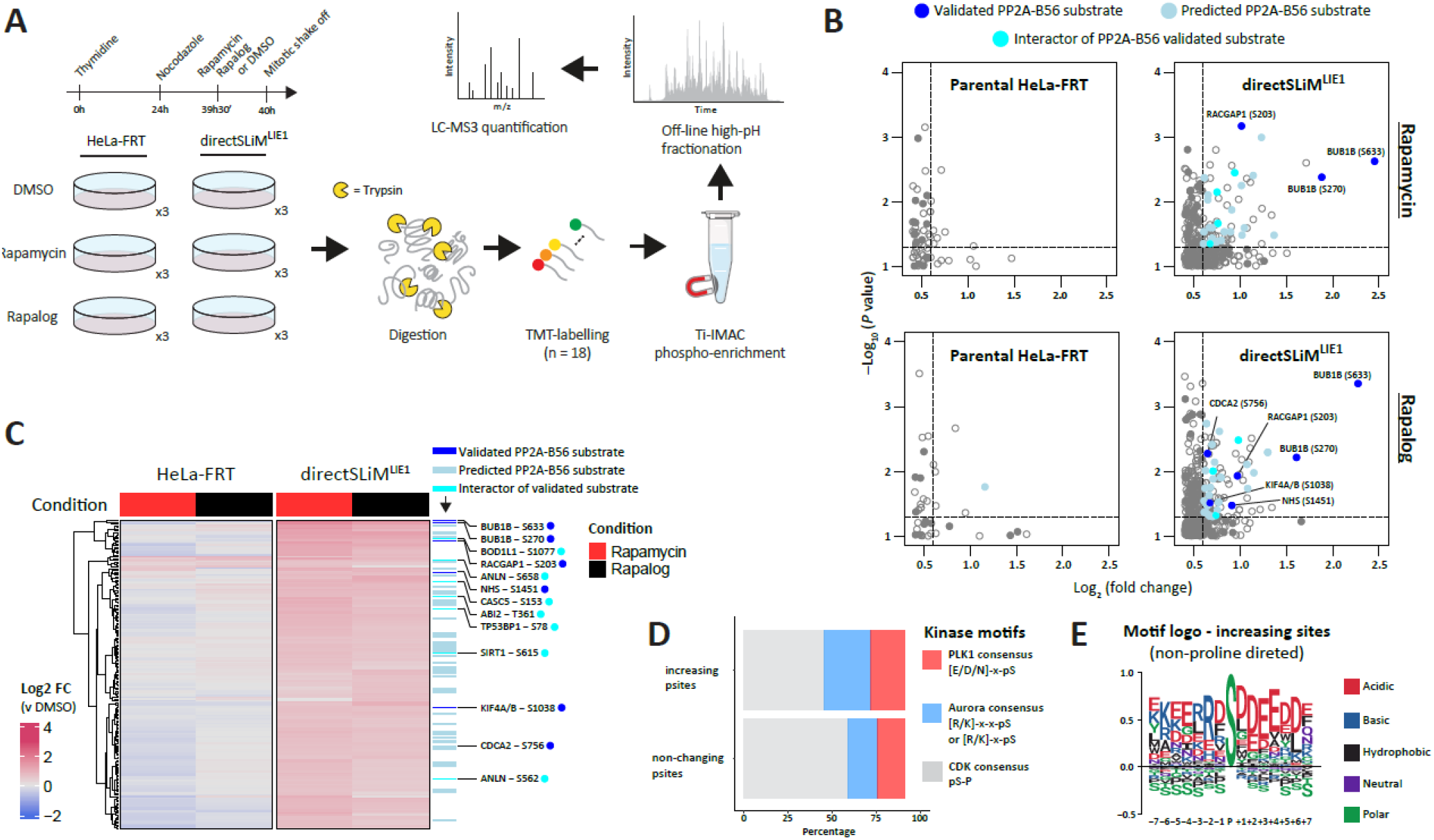
Global protein phosphorylation changes after PP2A-inhibition in directSLiM^LIE1^ cells. **A**. Experimental design: mitotically arrested LIE1^FRB^ and HeLa-FRT cells were treated with rapamycin, rapalog or vehicle before harvest. Following cell lysis, proteins were digested, and peptides were TMT-labeled. Next, phosphopeptides were enriched with Ti-IMAC magnetic beads, followed by high-pH reversed-phase fractionation and measurement by LC-MS. **B**. Zoomed-in right upper quadrant of volcano plot showing upregulated phosphorylation sites after treatment with rapamycin or rapalog. Solid dots are either validated substrates, predicted substrates or interactors of validated PP2A-B56 substrates (see methods for details). Only hits above a -Log_10_(P value = 0.05) and >1.5-fold change are color coded. **C**. Heat map of phosphorylation sites upregulated after PP2A-B56 inhibition in directSLiM^LIE1^ cells. Each row represents the intensity of a phosphorylation site (n = 187). Intensities are normalised to vehicle treated condition for each cell line to show the relative fold-change upon rapamycin/rapalog addition. Gene names are added for all phospho-sites from validated substrates or interactors of validates substrates **D**. Percentage of Aurora kinase, PLK1 and CDK consensus motifs in the increasing and non-changing phosphorylation sites. **E**. Sequence logo of non-proline directed upregulated phosphorylation sites (all non-changing phosphosites used as background).

There was good correlation between the different heterodimerisers in the directSLiM^LIE1^ expressing cell lines (Figure 2C, Supplementary Figure 1B), and similar changes were not observed in parental HeLa-FRT cells (Figure 2B-C), implying they were due to PP2A-B56 inhibition. There was also specific enrichment of validated (P = 0.004, OR = 5.1) and predicted (p = 0.03, OR = 1.5) LxxIxE-containing motifs within proteins that increased in phosphorylation upon drug treatment (Supplementary Figure 1C), and the most significant hits were on the well-characterised substrate BUBR1 (Figure 2B-C). The increasing phospho-sites were enriched for Aurora and PLK1 consensus motifs and depleted for proline-directed CDK motifs (Figure 2D-E). This is consistent with previous data showing that PP2A-B56 displays reduced activity against CDK substrates (25), and is mainly involved in antagonising PLK1 (17, 18), MPS1 (21) and Aurora B activities (15, 32) (note that MPS1 has a very similar substrate preferences to PLK1 (33, 34)).

To validate if the increase in B56 substrate phosphorylation can be observed using a complementary approach, we used antibodies against a wide range of phosphorylation sites on BUBR1 interacting proteins. BUBR1 binds directly to BUB1, and the BUB1:BUBR1 complex is recruited to kinetochores by binding to phosphorylated MELT motifs on KNL1 (13). The schematic in Figure 3A shows the complex along with relevant phospho-sites for which validated phospho-antibodies are available. Importantly, all of these phosphorylation sites are known to be regulated directly or indirectly by PP2A-B56 (15, 17, 18, 20, 35). Immunofluorescence analysis demonstrated the expected phosphorylation increases following rapamycin treatment in directSLiM^LIE1^ cells, implying localised inhibition of B56 at the kinetochore (Figure 3B and Supplementary Figure 2). However, we also noticed an increase in basal phosphorylation of these sites in the absence of rapamycin treatment, compared to untreated HeLa-FRT controls.

**Figure 3.**
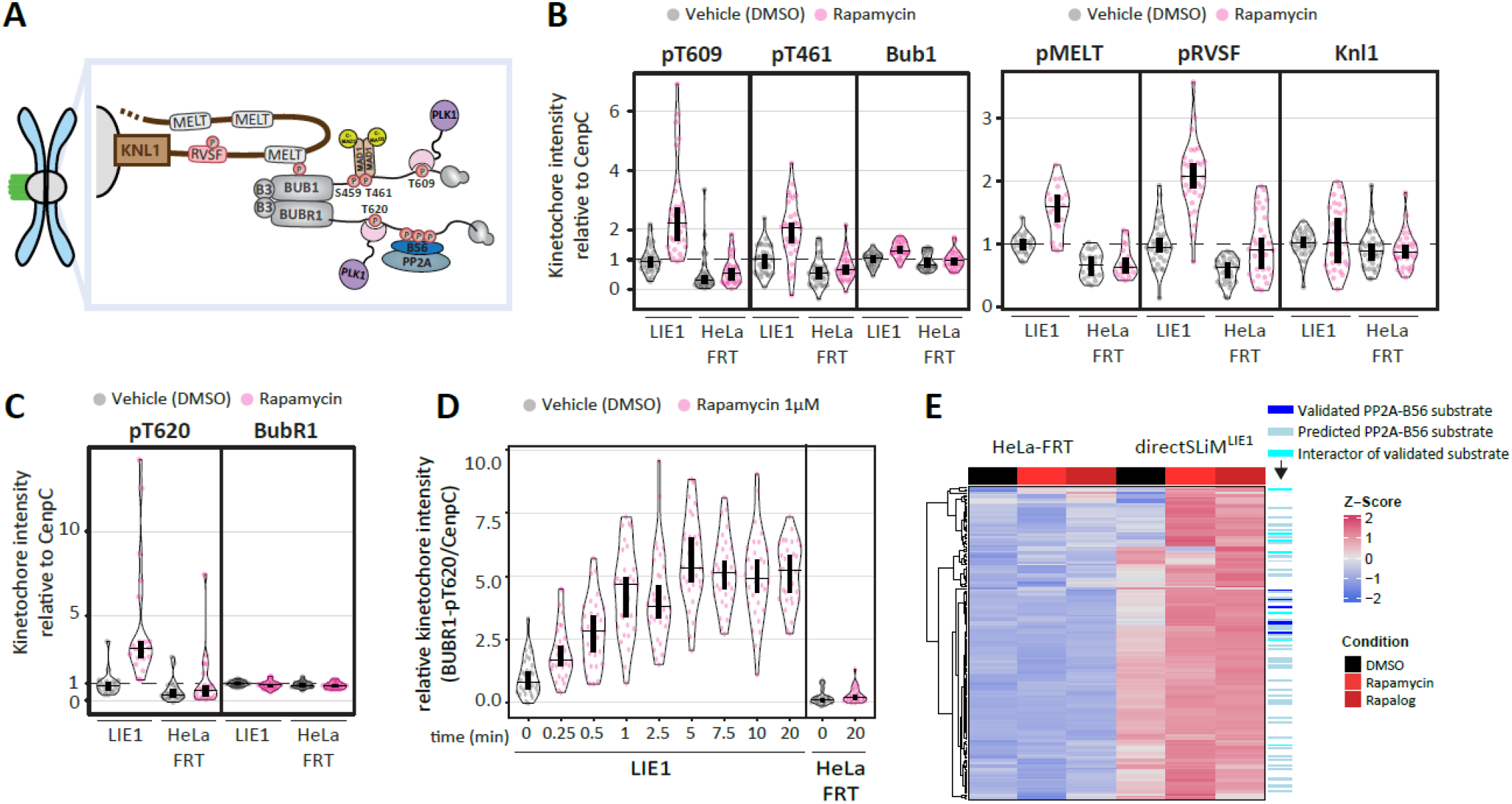
Basal inhibition of PP2A-B56 is observed in directSLiM^LIE1^ cells. **A**. Schematic illustrating kinetochore phosphosites on the BUB complex and KNL1 that are regulated by PP2A-B56. **B-C**. Effects of PP2A-B56 inhibition on the levels of BUB1-pT609, BUB1-pT461 and BUB1 (B, left) or KNL1-pMELT, KNL1-pRVSF and KNL1 (B, right) or BUBR1-pT620 and BUBR1 (C) at unattached kinetochores, in nocodazole-arrested HeLa FRT cells with/without directSLiM^LIE1^ expression and treated with vehicle or rapamycin for 20min. Kinetochore intensities from 20-30 cells, 2-3 experiments. **D**. Levels of BubR1-pT620 at unattached kinetochores in nocodazole-arrested directSLiM^LIE1^ cells treated with rapamycin for indicated times. Kinetochore intensities from 30 cells, 3 experiments **E**. Heat map of phosphorylation sites upregulated after PP2A-B56 inhibition in directSLiM^LIE1^ cells. Each row shows the z-scored intensity of a phosphorylation site (n = 170). Data information: Kinetochore intensities in B-C are normalized to LIE1 vehicle condition. Violin plots show the distributions of kinetochore intensities between cells. For each violin plot, each dot represents an individual cell, the horizontal line represents the median and the vertical one the 95% CI of the median, which can be used for statistical comparison of different conditions (see Methods).

We were initially confused as to why BUBR1-pT620, which is the closest to the B56 binding site (Figure 3A), did not also increase basally in directSLiM^LIE1^ cells (Figure 1E-G). However, this analysis was performed with a 1-min pre-extraction procedure to reduce background cytoplasmic staining and allow visualisation of R1A localisation to kinetochores. Repeat analysis of BUBR1-pT620 without pre-extraction showed increased basal phosphorylation in the absence of rapamycin treatment in directSLiM^LIE1^ cells (Figure 3C and Supplementary Figure 2). The pre-extraction may have also altered the apparent kinetics of inhibition because BUBR1-pT620 was observed to increase within 15 seconds of rapamycin-treatment without pre-extracting cells (Figure 3D).

We next wanted to assess whether basal inhibition was also apparent in our proteomic analysis. It is possible that substrates displaying increased basal phosphorylation in directSLiM^LIE1^ cells could be missed using a 1.5-fold cutoff upon rapamycin/rapalog treatment. Therefore, we filtered based on a >1.5-fold increase from DMSO-treated HeLa-FRT cells instead, and applied a less stringent requirement for a significant increase from DMSO-treated directSLiM^LIE1^ cells. This reanalysis showed that the sites that increased upon rapamycin/rapalog treatment in direct SLiM^LIE1^ cells had a tendency to have a higher level of basal phosphorylation compared to HeLa-FRT cells (Figure 3E), consistent with the basal inhibition observed in our immunofluorescence analysis. Together, this data implies that LIE1-B56 affinity is sufficiently strong to compete off some B56 interactors even in the absence of rapamycin. Therefore, we set out to generate new LIE sequences that had reduced affinity for B56, with the expectation that these should still be inhibitory when tethered to PP2A by heterodimeriser treatment.

We first used a range of LxxIxE peptides that had published Kd’s for B56 ranging from 750nM to 110uM, which was significantly higher than the 80nM Kd of LIE1 (Figure 4A) (7, 8, 12). The strongest binding peptide in these studies (LIE2) behaved similarly to LIE1, as assessed by BUBR1-pT620 immunofluorescence or live-cell imaging experiments to quantify chromosome alignment phenotypes that result from decreased BUBR1-B56 interaction (Figures 4B and Supplementary Figure 3A). Peptides with lower affinities than LIE2 (LIE3-7) exhibited reduced inhibition both in the absence or presence of rapamycin. Therefore, we next tried to remove key acidic residues C-terminal to the LxxIxE motif in the original LIE1 peptide, which is predicted to progressively decrease binding strength (7, 11, 12), creating 8 further LIE peptides (LIE8-14; Figure 4C). This identified that the sequence LIE9 was able to achieve strong B56 inhibition upon rapamycin treatment, but with minimal effects on chromosome alignment or BUBR1-T620 phosphorylation in the absence of rapamycin (Figures 4E and Supplementary Figure 3B). DirectSLiM^LIE10^ cells also had low basal effects and good inducible inhibition, however mitotic duration was still slightly extended compared to directSLiM^AAA^ controls (Supplementary Figure 3C), which may be due to subtle basal inhibition. DirectSLiM^LIE9^ cells also consistently produce no or negligible basal effects against other sites on BUB1 and KNL1 (Figure 4E and Supplementary Figure 3D).

**Figure 4.**
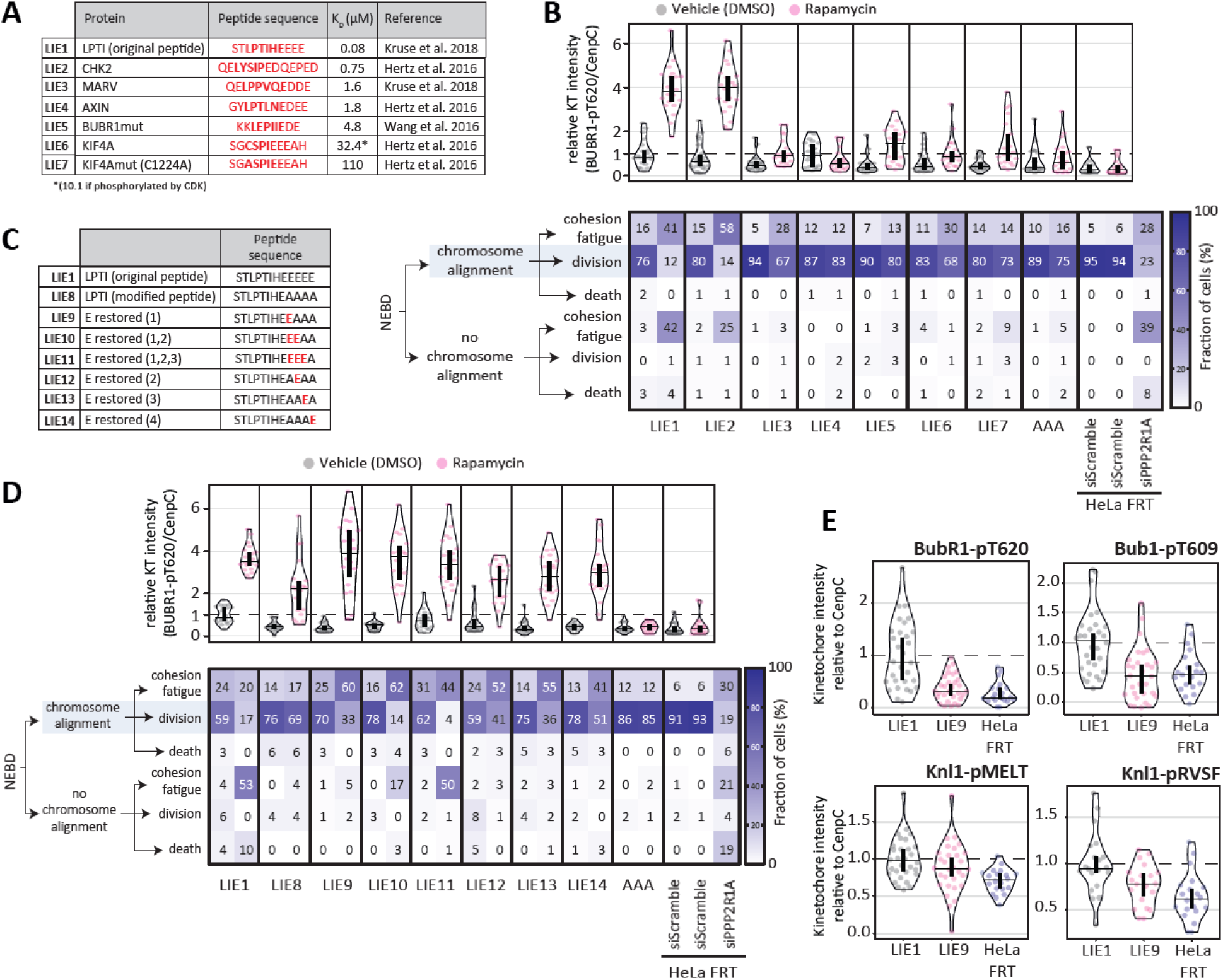
Optimization of the directSLiMs strategy to reduce basal inhibition of PP2A-B56. **A**. Table reporting the tested SLiMs with a lower binding affinity for PP2A-B56 in comparison to LIE1. **B**. Effects of PP2A-B56 inhibition on the levels of BUBR1-pT620 at unattached kinetochores (top) and on the mitotic cell fate after nuclear envelope breakdown (NEBD, bottom) in HeLa FRT cells with/without directSLiMs^LIE1-7/AAA^ expression and treated with vehicle or rapamycin. Top graph: Kinetochore intensities from 20 nocodazole-arrested cells, 2 experiments except HeLa-FRT, DMSO: 10 cells,. The treatment with vehicle/rapamycin was performed for 20min prior fixation. Bottom graph: heatmap showing the mean frequencies of cell fate after NEBD in each condition: 2 experiments, 17-50 cells per condition per experiment (see also Supplementary Figure 3A). **C**. Table reporting the tested SLiMs with different numbers of acid residues C-terminal to the LxxIxE motif. **D**. Effects of PP2A-B56 inhibition on the levels of BUBR1-pT620 at unattached kinetochores (top) and on the mitotic cell fate after nuclear envelope breakdown (NEBD, bottom) in HeLa FRT cells with/without directSLiMs^LIE1/LIE8-14/AAA^ expression and treated with vehicle or rapamycin. Top graph: Kinetochore intensities from 20 nocodazole-arrested cells, 2 experiments. The treatment with vehicle/rapamycin was performed for 20 min prior fixation. Bottom graph: heatmap showing the mean frequencies of cell fate after NEBD in each condition: 2 experiments, 42-50 cells per condition per experiment (see also Supplementary Figure 3B). **E**. Levels of BubR1-pT620, Bub1-pT609, Knl1-pMELT and Knl1-pRVSF at unattached kinetochores, in nocodazole-arrested HeLa FRT cells expressing directSLiMs^LIE1/LIE9^ and treated with vehicle for 20min. Note that the distributions of LIE1 and LIE9 are also shown in Supplementary Figure 3D. Kinetochore intensities from 20-30 cells, 2-3 experiments. Data information: Kinetochore intensities in B, D and E are normalized to LIE1 vehicle condition. Violin plots show the distributions of kinetochore intensities between cells. For each violin plot, each dot represents an individual cell, the horizontal line represents the median and the vertical one the 95% CI of the median, which can be used for statistical comparison of different conditions (see Methods).

We next performed new proteomic analysis in directSLiM^LIE9^ cells to identify phosphorylation changes following rapamycin or rapalog treatment. Considering rapamycin or rapalog treatment induced minimal phospho-changes in HeLa-FRT cells (Figures 2B-C), we decided to use directSLiM^AAA^ cells instead as the control condition this time, to examine if heterodimer-induced recruitment of FRB to PP2A-R1A impacted on PP2A function, for example, due to steric effects. We also used a modified phospho-peptide enrichment method that enhanced phospho-peptide recovery (Figure 5A, Supplementary Figure 4A). Using this method, we identified a total of 15,518 phospho-sites in mitotically arrest directSLiM^LIE9^ cells, of which 149 exhibited a significant >1.5-fold change following rapamycin or rapalog treatment. Some of the most significantly increasing phospho-sites were in well characterised PP2A-B56 substrates, including BUBR1 (BUB1B), RepoMan (CDCA2), RACGAP1, KIF4A and NHS (Figure 5B-C, Supplementary Figure 4B). There was also a strong significant enrichment for validated PP2A-B56 binding motifs in the proteins that contained increasing phospho-sites (adj.p = 3.1 ×10^−9^, OR = 12.5; Supplementary Figure 4C). We observed enrichment of potential Aurora kinase and PLK sites, and depletion of CDK sites (Figure 5D-E), as also observed in the previous MS analysis with directSLiM^LIE1^ cells (Figure 2D-E).

**Figure 5.**
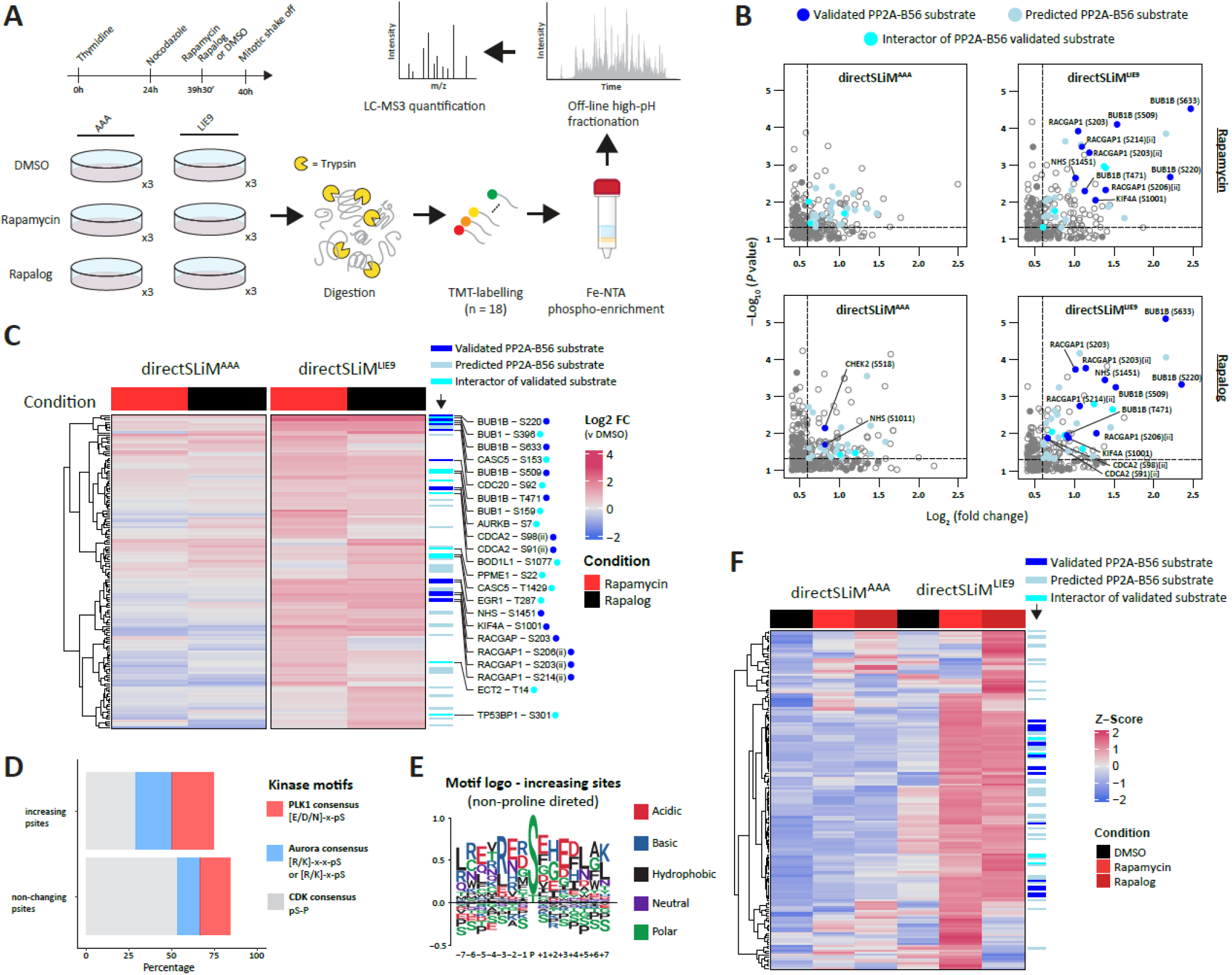
Global protein phosphorylation changes after PP2A-inhibition in directSLiM^LIE9^ cells. **A**. Experimental design: mitotically arrested LIE9^FRB^ and AAA^FRB^ cells were treated with rapamycin, rapalog or vehicle before harvest. Following cell lysis, proteins were digested, and peptides were TMT-labeled. Next, phosphopeptides were enriched with Fe-NTA spin column, followed by high-pH reversed-phase fractionation and measurement by LC-MS. **B**. Zoomed-in right upper quadrant of volcano plot showing upregulated phosphorylation sites after treatment with rapamycin or rapalog. Solid dots are either validated substrates, predicted substrates or interactors of validated PP2A-B56 substrates (see methods for details). Only hits above a -Log_10_(P value = 0.05) and >1.5-fold change are color coded. **C**. Heat map of phosphorylation sites upregulated after PP2A-B56 inhibition in directSLiM^LIE9^ cells. Each row represents the intensity of a phosphorylation site (n = 149). Intensities are normalised to vehicle treated condition for each cell line to show the relative fold-change upon rapamycin/rapalog addition. Gene names are added for all phospho-sites from validated substrates or interactors of validates substrates. **D**. Percentage of Aurora kinase, PLK1 and CDK consensus motifs in the increasing and non-changing phosphorylation sites. **E**. Sequence logo of non-proline directed upregulated phosphorylation sites (all non-changing phosphosites used as background). **F**. Heat map of phosphorylation sites upregulated after PP2A-B56 inhibition in directSLiM^LIE9^ cells. Each row represents the z-scored intensity of a phosphorylation site (n = 155). Label “(ii)” refers to phosphor-site in doubly phosphorylated peptides.

Basal phosphorylation levels were also generally lower in directSLiM^LIE9^ cells, when compared to the directSLiM^LIE1^ cells, using the same less stringent filtering we used to capture any basally high phosphorylation events (Figure 5F; compare to figure 3E). This indicates that LIE9-dependent PP2A-B56 inhibition is largely dependent on hetero-dimeriser treatment. We did observe more heterodimeriser-dependent changes in directSLiM^AAA^ cells than we had observed previously in parental HeLa-FRT cells (compared Figures 5B and 2B), but these were generally not the phospho-sites that increased further upon hetero-dimeriser treatment (Figure 5F). These increased phosphorylation sites could reflect steric effects of AAA^FRB^ recruitment to PP2A complexes. The correlation between the effects of rapamycin or rapalog was good, but not as high when compared to the directSLiM^LIE1^ cells (Figure 5C, Supplementary Figure 4D). This may be due to the lower affinity of the LIE9 sequence slowing the rate of PP2A-B56 inhibition, which could disproportionately affect rapalog-treated cells which were generally inhibited slower (Figure 1F-G). Importantly however, all increases to validated substrates, or to proteins that interact with validated substrates, increase following both rapamycin and rapalog treatment (Figure 5F). This suggests that the subset of targets that change with both drugs may be the most reliable.

The ability to rapidly inhibit PP2A-B56 within seconds (Figure 1F-G) creates opportunities to examine PP2A function in cell cycle phases that last only minutes. One good example is metaphase, which is the phase when all kinetochores have attached to microtubules. Metaphase last only 5-10 minutes but during that time the SAC signal is switched off, the mitotic checkpoint complex is disassembled, and active APC/C^CDC20^ degrades its key mitotic substrates cyclin B and securin, allowing cells to exit mitosis into anaphase (36). If kinetochores become detached during metaphase then the SAC is reimposed and mitotic progression is rapidly halted (37). PP2A-B56 is required to stably attach kinetochores to microtubules during prometaphase (9, 10, 38), but whether PP2A-B56 is still required to maintain those attachments on aligned chromosomes at metaphase remains unknown (Figure 6A). Fixed chromosome alignment assays in metaphase-arrested cells demonstrated that 30 min rapamycin treatment was sufficient to detach chromosomes from microtubules in directSLiM^LIE9^ cells, but not in directSLiM^AAA^ cells, which is consistent with reduced PP2A-B56 recruitment to BUBR1 inhibiting kinetochore-microtubule attachments (Figure 6B). However, PP2A-B56 also binds Sgo1 at the centromere to maintain centromeric cohesion, in a manner that is dependent on its LxxIxE-binding pocket (24). To test whether loss of centromeric cohesion contributes to the chromosome misalignment phenotypes at metaphase, we depleted WAPL because this can preserve cohesion in the absence of PP2A (39, 40). WAPL depletion was able to fully rescue misaligned chromosomes following Sgo1 depletion, as expected (39, 40), but it did not affect the detachment of chromosomes in directSLiM^LIE9^ cells ^treated^ with rapamycin (Figure 6B). Therefore, B56 inhibition at metaphase impairs chromosome alignment, most likely by dissociating PP2A from the outer kinetochore BUBR1 to detach kinetochore-microtubules.

**Figure 6.**
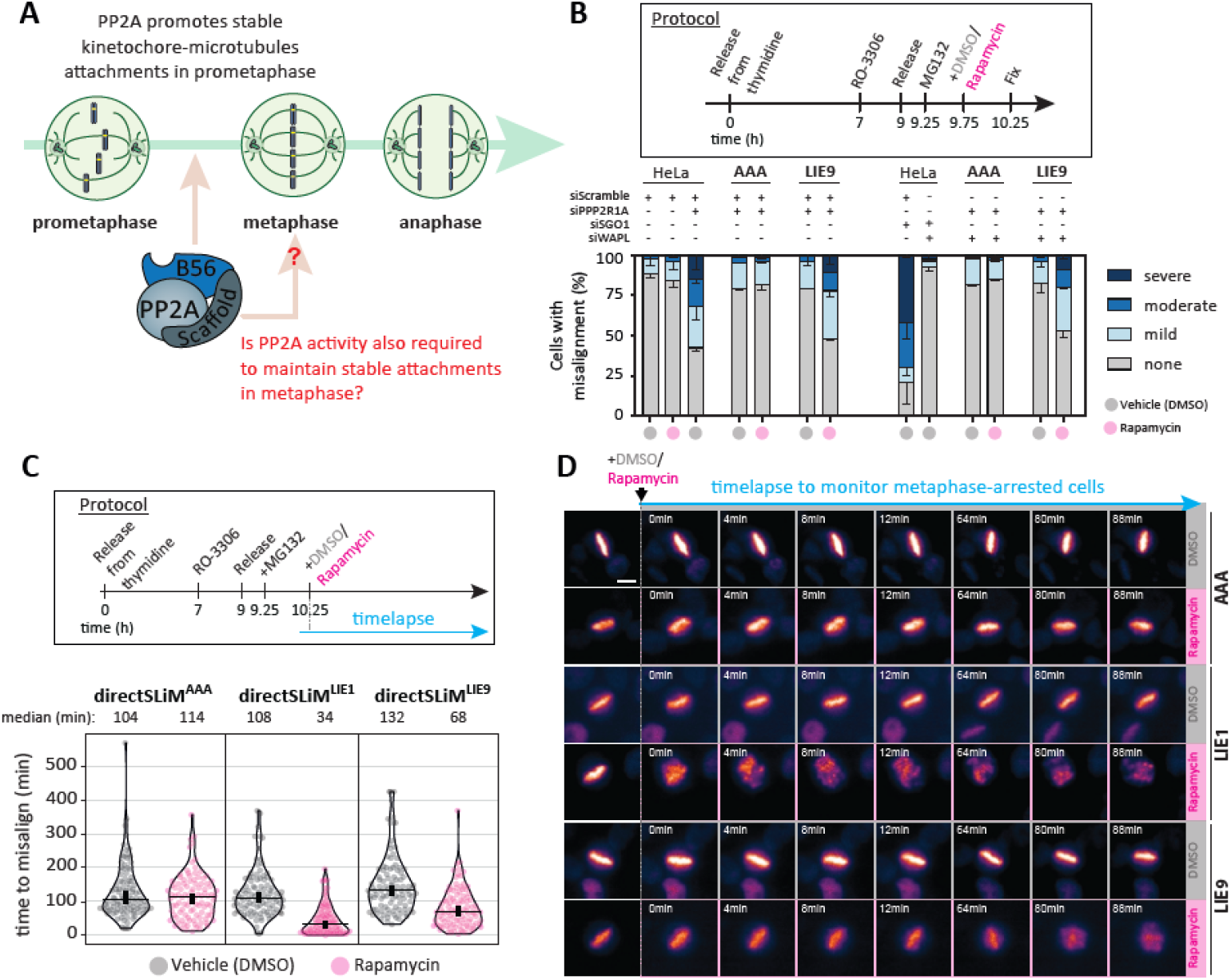
PP2A-B56 is needed to maintain stable kinetochore-microtubules attachment at metaphase. **A**. Schematic illustrating known and potential contributions of PP2A-B56 in stabilizing kinetochore-microtubule attachments during mitosis. **B**. Evaluating the effects on chromosome alignment in HeLa FRT cells with/without directSLiMs^LIE9/AAA^ expression and treated with vehicle or rapamycin. Top panel: protocol used to visualise chromosome alignment in fixed samples (see Materials and Methods for details). Bottom panel: graph showing mean frequencies of chromosome misalignments (±SEM) of 2 experiments, 100 cell quantified per condition per experiment. **C**. Evaluating the timings of chromosome misalignments during a metaphase arrest, in HeLa HistH1H4C-TagGFP2 cells expressing the directSLiMs^LIE1/LIE9/AAA^ and treated with vehicle or rapamycin. Top panel: protocol used to visualise chromosome misalignments in live cells (see Material and Methods for details). Bottom panel: timings of chromosome misalignments of 2 experiments, 50 cells quantified per condition per experiment. Violin plots show the distributions of misalignment timings between cells. For each violin plot, each dot represents an individual cell, the horizontal line represents the median and the vertical one the 95% CI of the median, which can be used for statistical comparison of different conditions (see Materials and Methods). **D**. example images of the misalignment timings shown in C (see Methods for details). Scale bar: 10 μm.

To determine the kinetics of kinetochore-microtubule detachment at metaphase, we used live assays to quantify the time to misalignment in metaphase-arrested directSLiM^LIE9^ cells treated with rapamycin. This revealed that a significant portion of cells lose their metaphase-aligned chromosomes during the first 30 min of rapamycin treatment, consistent with our fixed analysis (note that chromosome misalignment occurs in all metaphase arrested cells within approximately 2 hours due to cohesion fatigue (41)) (Figure 6C). Rapamycin treatment in metaphase-arrested directSLiM^LIE1^ cells, however, causes a more rapid and often immediate dissociation of chromosome from the metaphase plate (Figure 6C-D and supplementary movies 1-6). This rapamycin-dependent misalignment was also observed in fixed assays and was not caused by loss of centromeric cohesion because it was unaffected by WAPL depletion (Supplementary Figure 5). In summary, rapid PP2A-B56 inhibition can be used to uncover a critical role for PP2A-B56 in maintaining chromosome alignment at metaphase. This is most likely due to inhibition of PP2A-BUBR1 interaction leading to destabilised kinetochore-microtubule attachments.

## Discussion

Here we present a chemical-genetic system that can rapidly block the substrate binding pocket on PP2A-B56 thereby effectively inhibiting phosphatase activity. This is a powerful approach that can specifically inhibit PP2A-B56 function within seconds of drug addition, and we use this to identify PP2A-B56 substrates during mitosis and to characterise the role of PP2A-B56 during metaphase. It should now be possible to use this directSLiM system to carefully study PP2A-B56 function during any other cell cycle phase or biological process.

The directSLiM approach is also easily adaptable to inhibit other SLiM-binding interfaces on any other proteins. Care just needs to be taken to identify SLiMs with the optimal binding strength, so that inhibition is only observed upon drug addition. However, there may still be situations when strong binding SLiMs are preferred, for example, if strong inhibition or extremely rapid inhibition kinetics are required. A good example is the use of directSLiM^LIE1^ cells to cause immediate detachment of chromosomes at metaphase, which is not observed in directSLiM^LIE9^ cells (Figure 6C-D). Care also needs to be taken to assess the effects of drug alone, especially if rapamycin is used. In mitosis we observed very few phosphorylation changes within 30 minutes rapamycin treatment (Figure 2B, C), but this may be due to the fact that TORC1 activity is strongly inhibited during a mitotic arrest (42). Rapalog does not inhibit TORC1 and this also works well, albeit with slightly slower kinetics (Figure 1F-G). Other chemically-induced dimerisation systems could also be used, if required (43).

We use the directSLiMs system to demonstrate that PP2A-B56 is needed to maintain kinetochore-microtubule attachments at metaphase, in addition to its previous known role in stabilising initial kinetochore-microtubule attachments in prometaphase (9, 10, 38). This is important because it implies that PP2A-B56 is required for proper tension-sensing at kinetochores: a process that is still poorly understood (44). Aurora B activity is reduced at kinetochores that come under tension, thereby stabilising kinetochore-microtubule attachments on chromosomes that have aligned to the metaphase plate. The requirement for PP2A-B56 to maintain those stable attachments suggests that PP2A is also needed for kinetochores to respond correctly to tension-dependent reductions in Aurora B activity.

Our sequence analysis of the phosphorylated peptides that increase following rapamycin/rapalog treatment suggests PP2A-B56 is important to antagonise the mitotic kinases Aurora A/B, PLK1 and/or MPS1, as suggested by numerous previous studies (9, 10, 15, 17-21, 32, 38). Whether these phosphorylation sites increase because of increased kinase activity, decreased substrate dephosphorylation, or a combination of both, remains to be determined.

In summary, we present here a new chemical-genetic system to inhibit and characterise PP2A-B56, revealing new roles for PP2A in promoting stable kinetochore-microtubule attachments at metaphase. This system should be expandable to inhibit other phosphatase complexes that also rely on SLiM-based substrate recognition (3). The directSLiMs approach may therefore ultimately prove as valuable for the characterising phosphatases as the “bump-and-hole” approach has proved for characterising kinases that lack small molecules inhibitors (45). In fact, it could be more widely applied to characterise many other enzymes or proteins that are regulated by SLiM-based interactions. The eukaryotic linear motifs resource currently lists hundreds of annotated SLiM classes, regulating a wide range of processes including protein transcription and translation, cellular trafficking, cell cycle, and protein degradation (26). DirectSLiMs could therefore prove to be a powerful approach to dissect the role of these different SLiM classes once the SLiM-binding domains have been identified and validated.

## Supporting information

Supplementary Movie S6 - LIE9_Rapa

Supplementary Movie S1 - AAA_DMSO

Supplementary Movie S2 - AAA_Rapa

Supplementary Movie S3 - LIE1_DMSO

Supplementary Movie S4 - LIE1_Rapa

Supplementary Movie S5 - LIE9_DMSO

## Acknowledgements

We would like to thank the MRC PPU cloning service, MRC PPU sequencing service, and the light microscopy facility at Dundee. We also thank Geert Kops and Matthieu Bollen for reagents. This work was supported by a Wellcome Investigator grant to A.T.S that funds L.A., A.C. and J.M.V (222494/Z/21/Z).

## Author Contributions

A.T.S. conceived the project and supervised the study. L.A.A. developed the directSLiM system with help from A.C. J.M.V performed the MS experiments and analysis. A.C. performed the phenotypic chromosome alignment and segregation experiments. R.T. performed all cloning. T.L. co-supervised J.M.V. for the MS studies. A.T.S. wrote the initial draft of the paper, and all other authors generated figures and edited the paper.

## Materials and Methods

### Cell culture and reagents

All cell lines used in this study were derived from HeLa Flp-in cells (a gift from S Taylor, University of Manchester, UK) (46) apart from Phoenix Ampho 293 cells (ATCC). Cells were authenticated by STR profiling (Eurofins). Cells were cultured in full growth media – DMEM supplemented with 9% FBS and 50 µg/ml penicillin/streptomycin. Every 4-8 weeks, cells were screened to ensure a mycoplasma free culture. The following drugs were used (at the indicated concentrations throughout): Doxycycline (1μg/ml), thymidine (2mM), nocodazole (3.3μM) and MG132 (10μM) were purchased from Sigma Aldrich; rapamycin (1 μM) from LC Labs; A/C heterodimerizer (or Rapalog, 500nM) from Clontech; puromycin (1μg/ml) and hygromycin B (200μg/ml) from Santa Cruz Biotechnology; RO-3306 (10μM) from Tocris; the SiR-DNA far-red DNA probe (1:10000) from Spirochrome.

### Plasmids and cloning

All plasmids were cloned by MRCPPU Reagents & Services Laboratory. The DUXXXXX numbers indicated below relate to the plasmid numbers that can be ordered from MRC PPU Reagents (https://mrcppureagents.dundee.ac.uk/) To clone the directSLiM construct pcDNA5-LIE1-VSV-FRB-T2A-FKBP-3xFLAG-PPP2R1A (named LIE1^FRB^-T2A-^FKBP^R1A hereafter, with LIE1 sequence STLPTIHEEEEE), synthesised fragments for siRNA-resistant PPP2R1A and for LIE1-VSV-FRB-T2A-FKBP (Genestrings, Thermofisher) were subcloned into pcDNA5/FRT/TO using KpnI-BamHI and BamHI-ApaI respectively. To generate AAA^FRB^-T2A-^FKBP^R1A, the LPTI sequence in the LIE1 feature of LIE1^FRB^-T2A-^FKBP^R1A was replaced with AAAA by site-directed mutagenesis (STAAAAHEEEEE - DU71843, MRCPPU Reagents & Services Laboratory - https://mrcppureagents.dundee.ac.uk/). To generate LIE2-7^FRB^-T2A-^FKBP^R1A constructs, the LIE1 feature in LIE1^FRB^-T2A-^FKBP^R1A was replaced with the sequence for LIE2 (QELYSIPEDQEPED, DU75900), LIE3 (QELPPVQEDDE, DU75903), LIE4 (GYLPTLNEDEE, DU75898), LIE5 (KKLEPIIEDE, DU75904), LIE6 (SGCSPIEEEAH, DU75901) or LIE7 (SGASPIEEEAH, DU75902) by annealing the appropriate oligo pair and ligating into LIE1^FRB^-T2A-^FKBP^R1A digested with KpnI and AvrII (MRCPPU Reagents & Services Laboratory - https://mrcppureagents.dundee.ac.uk/). To generate LIE8^FRB^-T2A-^FKBP^R1A, the LIE1 feature of LIE1^FRB^-T2A-^FKBP^R1A was replaced with STLPTIHEAAAA by site-directed mutagenesis (DU75852, MRCPPU Reagents & Services Laboratory - https://mrcppureagents.dundee.ac.uk/). Site-directed mutagenesis with specific primers (Sigma-Alrich) was performed on LIE8^FRB^-T2A-^FKBP^R1A to generate: LIE9^FRB^-T2A-^FKBP^R1A (STLPTIHEEAAA; forward: 5’-GCCCACAATTCATGAAGAAGCAGCGGCAGGAGG-3’, reverse: 5’-CCTCCTGCCGCTGCTTCTTCATGAATTGTGGGC-3’), LIE10^FRB^-T2A-^FKBP^R1A (STLPTIHEEEAA; forward; 5’-GCTGCCCACAATTCATGAAGAAGAGGCGGCAGGAGGAGG-3’, reverse: 5’-CCTCCTCCTGCCGCCTCTTCTTCATGAATTGTGGGCAGC-3’), LIE11^FRB^-T2A-^FKBP^R1A (STLPTIHEEEEA; forward: 5’-GAAGAAGAGGAAGCAGGAGGAGGTTCCGG-3’, reverse: 5’-CCGGAACCTCCTCCTGCTTCCTCTTCTTC-3’), LIE12^FRB^-T2A-^FKBP^R1A (STLPTIHEAEAA; forward: 5’-CAATTCATGAAGCTGAGGCGGCAGGAGGAGGTTCC-3’, reverse: 5’-GGAACCTCCTCCTGCCGCCTCAGCTTCATGAATTG-3’), LIE13^FRB^-T2A-^FKBP^R1A (STLPTIHEAAEA; forward: 5’-CATGAAGCTGCAGAGGCAGGAGGAGGTTCC-3’, reverse: 5’-GGAACCTCCTCCTGCCTCTGCAGCTTCATG-3’) and LIE14^FRB^-T2A-^FKBP^R1A (STLPTIHEAAAE; forward: 5’-GAAGCTGCAGCGGAAGGAGGAGGTTCCGG-3’, reverse: 5’-CCGGAACCTCCTCCTTCCGCTGCAGCTTC-3’). All final plasmids were verified by sequencing carried out by MRCPPU Reagents and Services Sequencing Laboratory (https://dnaseq.co.uk/). To generate doxycycline-inducible PPP2R1A shRNAs we used the pSuperior system (OligoEngine) to create a pSuperior-shPPP2R1A-33 plasmid (pSP-shR1A-33). Oligos containing the shRNA sequence (forward: 5’-GATCCCCTTTTCCACTAGCTTCTTCATTCAAGAGATGAAGAAGCTAGTGGAAAATTTTTA and reverse: 3’-AGCTTAAAAATTTTCCACTAGCTTCTTCATCTCTTGAATGAAGAAGCTAGTGGAAAAGGG) were annealed and ligated into pSuperior Retro Puro (OligoEngine) using HindIII and BglII. The CRISPaint constructs pCas9-mCherry-Frame1, pCas9-mCherry-HistH1H4C, and pCRISPaint-TagGFP2-PuroR were purchased from Addgene (https://pubmed.ncbi.nlm.nih.gov/27465542/).

### Gene expression

HeLa Flp-in cells were stably generated to allow doxycycline-inducible expression of all the directSLiMs constructs. Cells were transfected with the relevant pcDNA5/FRT/TO directSLiMs plasmid and the Flp recombinase pOG44 (Thermo Fisher) using Fugene HD (Promega) according to the manufacturer’s instructions. Subsequently, stable integrants at the FRT locus were selected using hygromycin B for at least 2 weeks.

### Generation of HeLa pSP-PPP2R1A cell lines

Phoenix Ampho 293 cells (ATCC) were transfected with pSP-shR1A-33 using Fugene HD (Promega) according to the manufacturer’s instructions. After 24h, cells were washed and placed in fresh full growth media. Forty-eight hours after transfection, virus containing media was harvested and filtered using a 0.45uM filter before adding to HeLa-Flp-in cells. Cells were infected with virus 3 times before the media was replaced with full growth media for 48h. Cells were then selected with puromycin for 3 days. HeLa pSP-PPP2R1A cells were then used to stably express doxycycline-inducible directSLiMs constructs by following the same procedure described above.

### Generation of HistH1H4C-tagged cell lines

The endogenous HistH1H4C in HeLa Flp-in cells was tagged with TagGFP2 by using the CRISPaint method (47). Briefly, cells were transiently transfected with pCas9-mCherry-Frame1 frame selector, pCas9-mCherry-HistH1H4C target selector and pCRISPaint-TagGFP2-PuroR donor plasmid at a ratio of respectively 1:1:2, using Fugene HD (Promega) according to the manufacturer’s instructions. After 2 days from transfection, cells were selected using puromycin for 2 weeks. These cells were then used to stably express doxycycline-inducible directSLiMs^LIE1/LIE9/AAA^ constructs by following the same procedure described above.

### Gene knockdown

For all experiments involving expression of the directSLiMs constructs in HeLa Flp-in or HeLa pSP-PPP2R1A cells, the endogenous mRNA of PPP2R1A was knocked down (siPPP2R1A: 5ʹ-CCACCAAGCACAUGCUACC-3’, dTdT overhang) and replaced with an siRNA-resistant FKBP-FLAG-tagged mutant expressed from the doxycycline-inducible directSLiMs construct. The other siRNAs used in this study were: siSGO1 (5ʹ-GAUGACAGCUCCAGAAAUU-3’, UU overhang), siWAPL (5’-GAGAGAUGUUUACGAGUUU-3’, UU overhang) and siScramble (control siRNA: 5ʹ-AAGCGCGCTTTGTAGGATTCG-3’, UU overhang). All synthesised siRNAs (Sigma-Aldrich) were used at 20nM final concentration. All siRNAs were transfected using Lipofectamine® RNAiMAX Transfection Reagent (Thermo Fisher) according to the manufacturer’s instructions. After 16 h of knockdown, cells were arrested with thymidine for 24 h. Doxycycline was used to induce the expression of the directSLiMs constructs – and to express PPP2R1A shRNA construct in HeLa pSP-PPP2R1A cells - during and following the thymidine block. Cells were then released from thymidine block into full growth media supplemented with doxycycline and, when appropriate, nocodazole for 5-7 hours for live imaging or 8.5 hours before processing for fixed analysis. Data was obtained using the following cell lines expressing directSLiMs: HeLa Flp-In (Fig 1 & Fig 6C-D); HeLa HistH1H4C-TagGFP2 cells (Fig 7C-D); HeLa Flp-In pSP-PPP2R1A cells (all other figures).

### Immunofluorescence

Cells plated on High Precision 1.5H 12-mm coverslips (Marienfeld) were fixed with 4% paraformaldehyde (PFA) in PBS for 10 min or pre-extracted with 0.1% Triton X-100 in PEM (100 mM PIPES, pH 6.8, 1 mM MgCl_2_ and 5 mM EGTA) for 1 minute before addition of 4% PFA for 10 minutes. Pre-extraction was only performed in cells shown in Figure 1E-G. After fixation, coverslips were washed with PBS and blocked with 3% BSA in PBS + 0.5% Triton X-100 for 30 min, incubated with primary antibodies overnight at 4°C, washed with PBS and incubated with secondary antibodies plus DAPI (4,6-diamidino2-phenylindole, Thermo Fisher) for an additional 2-4 hours at room temperature in the dark. Coverslips were washed with PBS and mounted on glass slides using ProLong antifade reagent (Molecular Probes). All images were acquired on a DeltaVision Core or Elite system equipped with a heated 37°C chamber, using a CoolSNAP HQ or HQ2 camera (Photometrics) with a 100x/1.40 NA U Plan S Apochromat objective using softWoRx software (Applied precision), or on a Nikon Ti2-E Eclipse system equipped with a heated 37°C Okolab chamber, using a Kinetix camera (Teledyne Photometrics) with a CFI Plan Apochromat λD 100x/1.45 NA oil objective (Nikon) with NIS-Elements AR software (Nikon). Images were acquired at 1×1 binning and processed using softWorx software, Nikon NIS Elements and ImageJ (National Institutes of Health). Mitotic cells arrested in early prometaphase were selected for imaging based on good expression of FLAG in the cytoplasm. All immunofluorescence images displayed are maximum intensity projections of deconvolved stacks and were chosen to closely represent the median quantified data. Figure panels were creating using Omero (http://openmicroscopy.org).

The following primary antibodies (all diluted in 3% BSA in PBS) were used at the final concentration indicated: guinea pig anti-CENP-C (PD030 from Caltag + Medsystems, 1:5000), rabbit anti-BUB1 (A300-373A from Bethyl, 1:1000), rabbit anti-BUBR1 (A300-386A from Bethyl, 1:1000), rabbit anti-KNL1 (ab70537 from abcam, 1:1000), mouse anti-FLAG(M2) (F3165-.2MG from Sigma, 1:1000).

The rabbit anti-pMELT-KNL1 antibody is directed against phospho-Thr 943 and -Thr 1155 of human KNL1 (15) (1:1000 – gift from G. Kops, Hubrecht, NL). The rabbit anti-pRVSF-KNL1 was raised against phospho-Ser 60 of human KNL1 (using the peptide C-CKKNSRRV[pS]FADTIK, custom raised by Biomatik, 1:500). The rabbit anti-BUBR1-pT620 antibody was raised against phospho-Thr 620 of human BUBR1 using the peptide C-AARFVS[pT]PFHE (custom raised by Moravian, 1:1000) (17). The rabbit anti-BUB1-pT609 antibody was raised against phospho-Thr 609 of human BUB1 using the peptide C-AQLAS[pT]PFHKLPVES (custom raised by Biomatik, 1:2000) (17). The rabbit anti-BUB1-pT461 against phospho-Thr 461 of human BUB1 was a gift from M. Bollen (Leuven, BE; 1:500) (35).

Secondary antibodies used were highly-cross absorbed goat anti-chicken Alexa Fluor 488 (A-11039), goat anti-rabbit Alexa Fluor 568 (A-11036), goat anti-mouse Alexa Fluor 488 (A-11029), goat anti-mouse Alexa Fluor 568 (A-11031), goat anti-guinea pig Alexa Fluor 647 (A-21450), donkey anti-rabbit Alexa Fluor 647 (A-31573) or donkey anti-mouse Alexa Fluor 647 (A-31571) all used at 1:1000 (Thermo Fisher).

### Immunoprecipitation and Western blotting

Endogenous PPP2R1A was knocked down with siRNA as above. Doxycycline was added 1h after siRNA transfection. After 24h, cells were treated with thymidine and doxycycline for 24 hr and then released into fresh media supplemented with nocodazole and doxycycline for 17 hr. After addition of DMSO or Rapamycin as indicated, mitotic cells were isolated by mitotic shake off, washed with ice-cold PBS and lysed in lysis buffer [50 mM Tris, pH 7.5, 150 mM NaCl, 1mM EDTA, 1% TX-100, phosStop and complete protease inhibitor cocktail, EDTA-free (both Roche)] on ice. Lysates were then incubated at 4°C on a rotating wheel for 20 min before centrifugation to remove insoluble material. 2-3mg lysate was incubated with 20µl FLAG-M2 magnetic beads (Sigma) for 3 hr at 4°C on a rotating wheel. The beads were washed 3x with wash buffer [50 mM Tris, pH 7.5, 150 mM NaCl, phosStop and complete protease inhibitor cocktail containing EDTA (both Roche)]. The sample was eluted by boiling the beads in SDS-PAGE gel loading buffer (62.5mM Tris, 2.5% SDS, 10% glycerol and 5% 2-mercaptoethanol) for 5min. Samples were processed for SDS-PAGE and immunoblotted using standard protocols.

The following primary antibodies (all diluted in 5% non-fat milk in TBST) were used at the final concentration indicated: mouse anti-B56a (BD Biosciences 610615, 1:2000), mouse anti-B56g (Santa Cruz Biotechnology sc-374379, 1:1000), mouse anti-B56d (Santa Cruz Biotechnology sc-271363, 1:1000), rabbit B56e (Aviva ARP56694-P050, 1:1000), mouse anti-PPP2CA (EMD Millipore 05-421, 1:1000), rabbit anti-PPP2R1A (Genetex GTX102206, 1:1000), rabbit anti-GEF-H1 (Abcam 155785, 1:1000), rabbit anti-BubR1 (Bethyl Laboratories A300-386A, 1:1000), rabbit anti-CDCA2 (Repoman, Sigma HPA030049, 1:1000), mouse anti-FLAG (Sigma F1804, 1:5000) and rabbit anti-Actin (Sigma A2066, 1:2500). The secondary antibodies were goat anti-mouse IgG HRP conjugate (Bio-Rad 170–6516, 1:2000) and goat anti-rabbit IgG HRP conjugate (BioRad 170– 6515, 1:5000). The secondary antibody used to detect B56α, B56γ, B56δ and FLAG was goat anti-mouse light chain specific HRP-conjugated (Sigma, AP200P, 1:1000).

### Chromosome alignment assays

To observe live chromosome alignment and determine mitotic cell fates and timing, cells were plated in 8-well or 18-well chamber slides (ibidi), released from thymidine block for 5-6 hours, incubated with SiR-DNA far-red DNA probe (1:10000, Spirochrome; to prevent toxicity (48)) in full growth media for 15 min, and treated with DMSO or rapamycin prior to imaging. Images were captured, after treatment with DMSO or rapamycin, every 4 minutes for 16 hours with a CFI Plan Apochromat λD 40x/0.95 NA air objective (Nikon) using a Nikon Ti2-E Eclipse with a Kinetix camera (Teledyne Photometrics) at 4×4 binning, 10 z-stacks with a step size of 1.50 µm. Selected cells were scored based on the following mitotic events: cohesion fatigue, cell division or cell death following chromosome alignment or not.

To observe chromosome alignment in fixed-cell experiments, cells were released from thymidine block for 7 hours before being treated for 2 hours with RO-3306 to synchronise cells at the G2/M boundary. Cells were then washed three times and incubated for 15 minutes with full growth media before addition of MG132 to prevent mitotic exit. After 30’ from MG132 addition, cells were treated with DMSO or rapamycin and fixed 30’ after the treatment. Fixed cells were stained as described above and imaged on a Zeiss Axio Observer with a CMOS Orca flash 4.0 camera at 4×4 binning, using a Plan-apochromat 20×/0.4 air objective, or on a Nikon Ti2-E Eclipse with Kinetix camera (Teledyne Photometrics) at 4×4 binning, using a CFI Plan Apochromat λD 20x/0.8 NA air objective. Cells with good expression of FLAG-tagged PPP2R1A were scored based on the number of misaligned chromosomes as aligned (0 misaligned chromosomes, with a visible metaphase plate), mild (1-2), moderate (3-5) or severe (>6).

To determine chromosome misalignment timings during an arrest in metaphase, HeLa HistH1H4C-TagGFP2 cells were plated in 8-well slides (ibidi), released from thymidine block for 7 hours and treated with RO-3306 and MG132 as described above, to enrich metaphase-arrested cells. After 1h from MG132 treatment, cells were treated with DMSO or rapamycin. To monitor chromosome misalignments of those cells arrested in metaphase prior to DMSO/rapamycin treatment, the imaging started 30’ after MG132 treatment, paused during the DMSO or rapamycin treatment and then resumed. Images were captured every 4 minutes for 12 hours with a CFI Plan Apochromat λD 40x/0.95 NA air objective (Nikon) using a Nikon Ti2-E Eclipse with a Kinetix camera (Teledyne Photometrics) at 4×4 binning, 10 z-stacks with a step size of 1.50 µm.

### Image quantification and statistical analysis

For quantification of kinetochore protein levels, images of similarly stained experiments were acquired with identical illumination settings and analysed using an ImageJ macro, as described previously (49). Fluorescence intensities at kinetochores were normalised to CenpC (i.e. kinetochore marker). Violin plots were produced using PlotsOfData - https://huygens.science.uva.nl/PlotsOfData/ (50). This allows the spread of data to be accurately visualised along with the 95% confidence intervals (thick vertical bars) calculated around the median (thin horizontal lines). This representation allows the statistical comparison between all treatments and timepoints because when the vertical bar of one condition does not overlap with one in another condition the difference between the medians is statistically significant (p<0.05). Heatmaps showing mitotic cell fates after nuclear envelope breakdown were generated with Graphpad Prism 7.

### Phosphoproteomics: Cell culture and sample preparation

After 24 h of knockdown and doxycycline addition, directSLiM cells were arrested with thymidine for 24 h. Cells were then released from thymidine block into full growth media with doxycycline and nocodazole for 15 hours. Cells were treated with either DMSO (0.1%), Rapamycin or rapalog for 30 minutes before harvesting. Mitotic cells were collected in their own media by shake-off, followed by a quick wash in plain DMEM containing doxycycline and the respective drug. Cells were then lysed with 2% SDS supplemented with protease and phosphatase inhibitors and snap-frozen in liquid nitrogen.

Cell lysates were subsequently cleared by sonication and nucleic acids were digested with Benzonase for 30 min at 37°C. Proteins were precipitated with acetone and digested into peptides with trypsin (1:100 enzyme to protein ratio) for 16 h at 37°C, followed by a second round of digestion for 4 h at 37°C. Next, peptides were acidified with formic acid (final concentration of 3%) and desalted with C18 QuikPrep® Micro SpinColumns™ from Harvard Apparatus. Briefly, columns were conditioned with 100% acetonitrile and then equilibrated with 0.5% formic acid. Peptides were loaded into the columns and washed twice with 0.5% formic acid followed by peptide elution using 80% acetonitrile in 0.5% formic acid. Peptides were dried in a vacuum concentrator at 30°C. Next, each sample was dissolved in 100 mM TEAB, mixed with 0.25 mg of TMTpro label and incubated for 1 h at room temperature. Labelling reaction was quenched by adding 2.5 uL of 5% hydroxylamine to each sample and incubating for 15 minutes at 37°C. Next, all samples were pooled together and dried before C18 desalting using 50 mg Sep-Pak C18 columns. Briefly, the TMT-labeled peptides were loaded into the columns, followed by washes with 0.5% formic acid and elution with 80% acetonitrile in 0.5% acetic acid.

Depending on the experiment, two different methods for phosphopeptide enrichment were used. For the experiment in Figure 2, peptides were mixed with MagResyn Ti-IMAC HP in 80% acetonitrile, 5% trifluoroacetic acid (TFA), 5% glycolic acid and incubated for 20 min at 25°C. Beads were washed with 80% acetonitrile, 1% TFA for 2 min, followed by a second wash with 10% acetonitrile, 0.2% TFA. Phosphopeptides were eluted twice with 1% ammonium hydroxide for 15 min, followed by a final elution with 50% acetonitrile, 1% ammonium hydroxide for 1 h. To increase phosho-peptide recovery, the flowthrough was mixed with a new batch of MagResyn Ti-IMAC HP beads followed by the same steps previously described. For the experiments in Figure 5, phospho-peptides were enriched using the High-Select Fe-NTA Phosphopeptide Enrichment Kit using the manufacturer instructions. Briefly, peptides were reconstituted in Binding/Wash Buffer and loaded into the Fe-NTA columns with an incubation time of 30 min at room temperature, gently mixing the resin every 10 min. Next, the column was washed a total of three times with Binding/Wash Buffer, before a final wash with LC-MS grade water. Phosphopeptides were eluted with Elution Buffer and acidified to 5% formic acid.

To ensure removal of magnetic beads or resin, the phosphopeptides were dried and desalted using C18 QuikPrep® Micro SpinColumns™ from Harvard Apparatus as previously described. To achieve a deep coverage of the phosphoproteome, peptides were fractionated using high-pH reversed-phase chromatography. Briefly, peptides were injected into a 1.0 × 100 mm column packed with 1.7 µm BEH particles with a 130 Å pore size coated with C18. For elution, we used a 15-80% B gradient using the following mobile phases: A, was 10 mM ammonium formate pH 9.0 in water and B was 10% ammonium formate pH 90 and 90% acetonitrile. Peptides were eluted, dried and stored at -20°C until measurement by LC-MS.

### Phosphoproteomics: Data acquisition

The experiment in Figure 2 was measured using a Dionex Ultimate 3000 HPLC coupled to an Orbitrap Fusion Tribrid mass spectrometer. Peptides from 14 fractions were loaded and separated using 75 μm × 50 cm EASY-Spray column with 2 μm sized particles. The column was kept at 50°C using an EASY-Spray source. The following mobile phases were used for the gradient: Buffer A consisted of 0.1% formic acid in LC-MS grade water and buffer B consisted of 80% acetonitrile and 0.1% formic acid. With a flow rate of 300 µL/min a gradient was applied, starting from 5% to 35% B in 130 minutes, followed by a 20 min wash with 98% B and a subsequent column equilibration with 5% for 20 min. A voltage of 2.2 kV was set for electrospray ionization and a MS1 scan on the Orbitrap was acquired at a 120,000 resolution with a maximum injection time of 50 ms with a scan range of 380-1500 m/z. The cycle time (time between MS1 scans) was set to 3 s. For peptide identification, precursors ions with a charge state of 2-6 were isolated (0.7 m/z isolation window) for CID fragmentation at 35% and subsequent MS2 on the Orbitrap at a 30,000 resolution with a maximum injection time of 60 ms. Dynamic exclusion was set for a duration of 45 s. 10 precursor fragments were selected for synchronous precursor selection (SPS) using an isolation window of 0.7 m/z and subjected to HCD at 65% for release of the TMTpro reporter tag and measured in the Orbitrap at 60,000 resolution with a maximum injection time of 105 ms.

The experiments in Figures 5 were measured using a Vanquish Neo UHPLC coupled to an Orbitrap Eclipse Tribrid mass spectrometer. Peptides from 16 fractions were separated using a 75 μm × 50 cm EASY-Spray PepMap Neo column with 2 μm sized particles coated with C18. The column was kept at 50°C using an EASY-Spray source. The following mobile phases were used for the gradient: Buffer A consisted of 0.1% formic acid in LC-MS grade water and buffer B consisted of 80% acetonitrile and 0.1% formic acid. A flow rate of 300 µL/min was used to apply a gradient from 2% to 40% B in 150 min, followed by a wash with 95% B for 20 minutes. A voltage of 1.9 kV was set for electrospray ionization and an MS1 scan on the Orbitrap was acquired at a 120,000 resolution with a maximum injection time of 50 ms and a scan range of 380-1500 m/z. The cycle time (time between MS1 scans) was set to 3 s. For peptide identification, precursors with charge state of 2-7 were isolated (0.7 m/z isolation window) for HCD fragmentation at 28% and subsequent MS2 on the ion trap with a maximum injection time of 50 ms. Dynamic exclusion was set for a duration of 70 s. 5 precursor fragments were selected for SPS using an isolation window of 0.7 m/z and subjected to HCD at 55% for release of the TMTpro reporter tag, followed by measurement in the Orbitrap at 50,000 resolution with a maximum injection time of 90 ms.

### Phosphoproteomics: Data analysis

Raw data were processed using MaxQuant (version 2.4.9.0) with the default settings for a TMTpro-18plex experiment. Carbamidomethyl (C) was added as a fixed modification while Oxidation (M), Acetyl (Protein N-term) and Phospho (STY) were added as variable modifications with a maximum number of 5 modifications per peptide. Database search was done against the human Swiss-Prot database, accessed on October 2023. Phosphorylation site intensities from the pSTY table were loaded into Perseus (version 2.0.5.0) and filtered to remove contaminants, decoy sequences and to keep only Class I phosphosites (localization probability score >0.75). Data was log2 transformed, normalized by median subtraction for each sample and subjected to statistical testing (T-test). Data was subsequently loaded into R Studio (2024.04.1+748) for clustering and visualization using the EnhancedVolcano, ComplexHeatmap and ggplot2 packages. Fisher tests to assess enrichment of validated and predicted PP2A substrates were performed using the database reported by (16). Motif analysis was performed by using the sequence window of each phosphosite for detection of kinase motifs using regular expressions in R applying these rules:

− PLK: [D/N/E]-X-[S/T]*
− Aurora kinases: [K/R]-X-[S/T]*[^P] or [K/R]-X-X-[S/T]*[^P]
− Cdk minimal consensus motif: [S/T]*-P

Where [] groups multiple residues for one position, * indicates the phosphorylated position, ^ before a certain residue indicates that it is forbidden for that position, and X represents any amino acid. The sequence logo of increasing phosphorylation sites against a background of all non-changing phosphorylation sites was generated using ggseqlogo package in R.

