## Supplementary Figures for "A chemical-genetic system to rapidly inhibit the PP2A-B56 phosphatase"

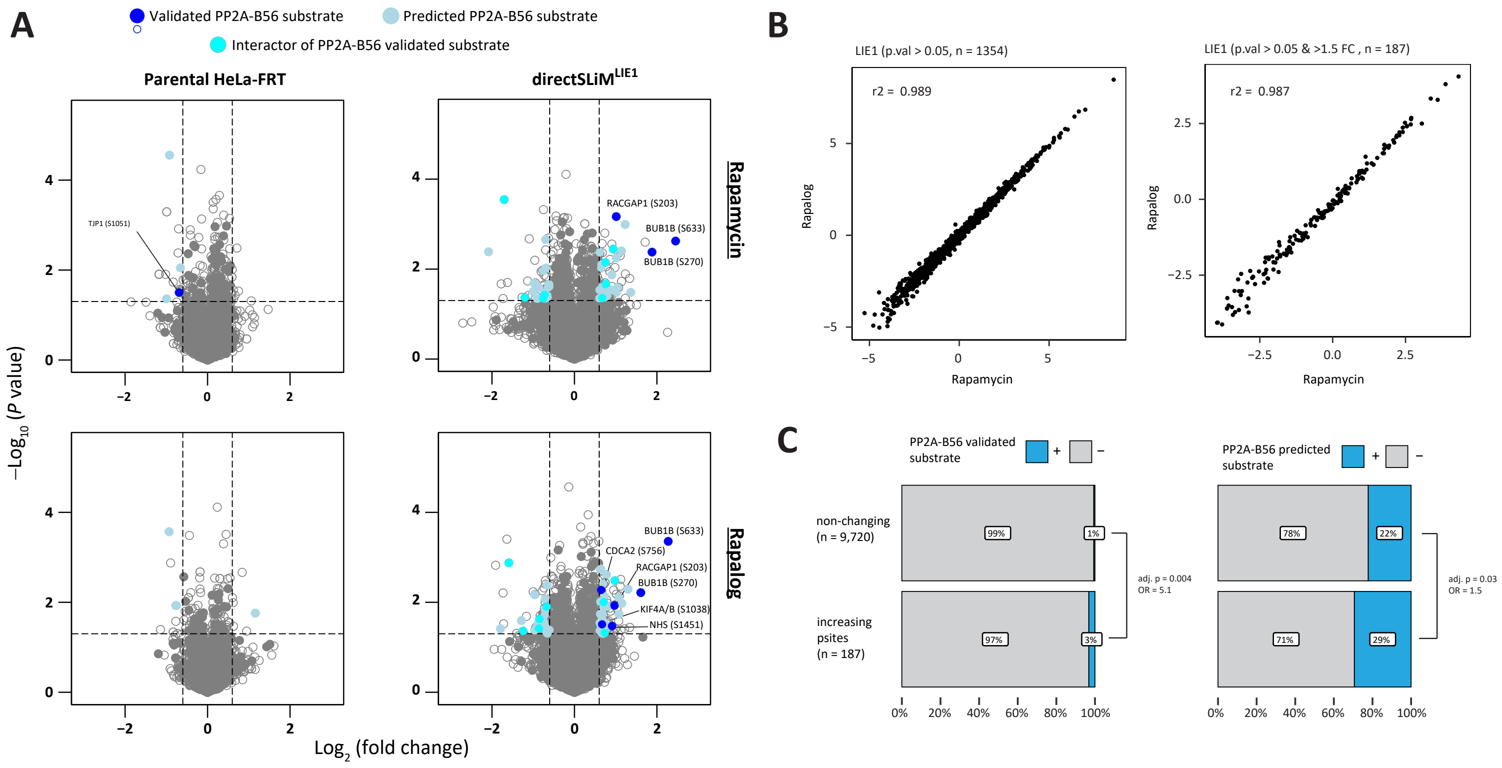


**Supplementary Figure 1 (related to Figure 2). A.** Volcano plot showing upregulated and downregulated phosphorylation sites after treatment with rapamycin or rapalog in directSLiM$^{LIE1}$ cells. Solid dots are either validated substrates, predicted substrates or interactors of validated PP2A-B56 substrates (see methods for details). Only hits above a -Log10(P value = 0.05) and >1.5-fold change are color coded. **B.** Scatter plot showing correlation of rapamycin and rapalog treatment in all changing sites (left) and in the top changing sites (right) in directSLiM$^{LIE1}$ cells. **C.** Assessing enrichment of PP2A-B56 validated (left) and predicted (right) substrates in proteins with increasing phosphorylation sites.

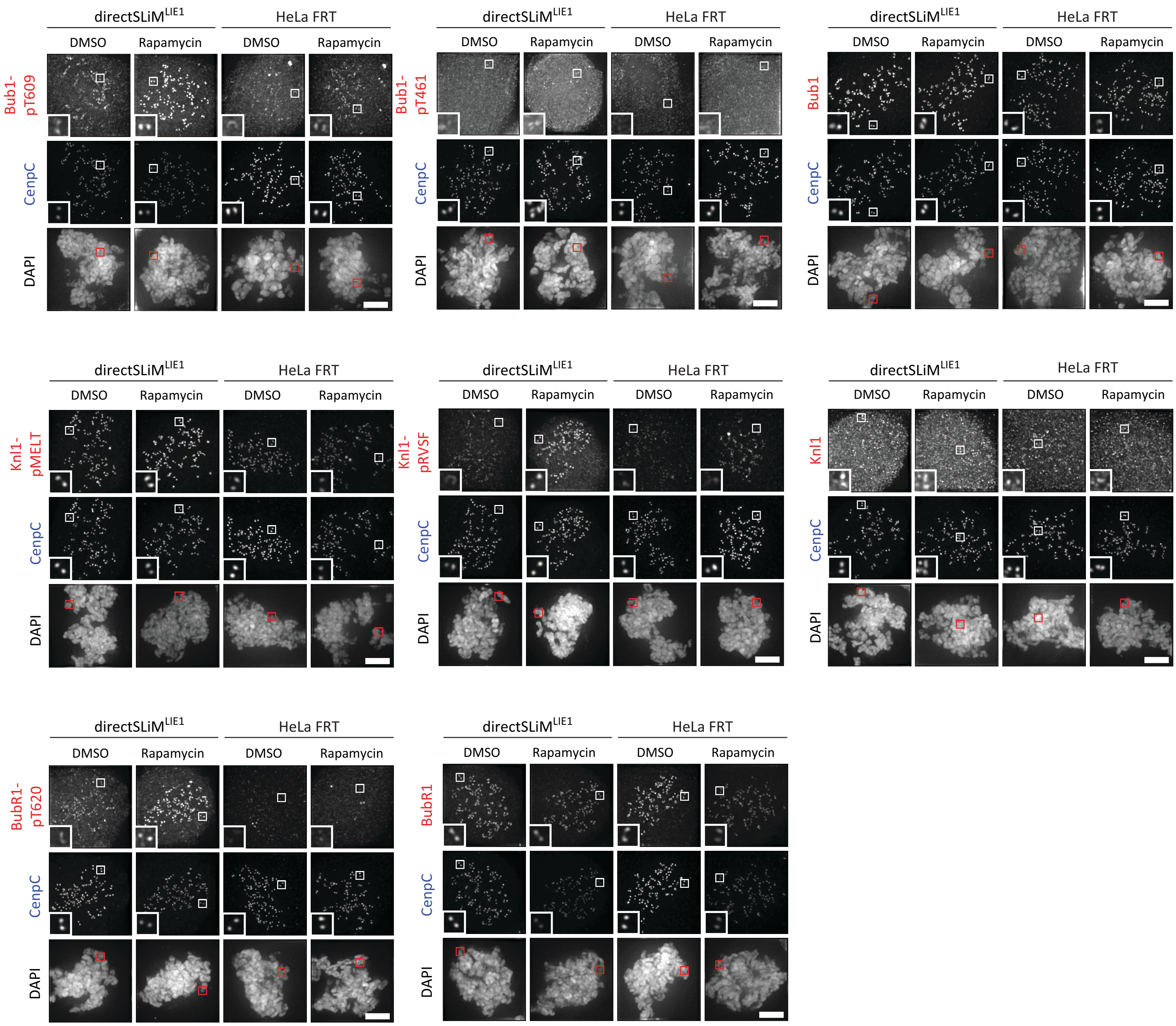


**Supplementary Figure 2 (related to Figure 3).** Representative example immunofluorescence images of the kinetochore quantifications shown in Figure 3B-C. The insets show magnifications of the outlined regions. Scale bars: 5µm. Inset size: 1.5µm

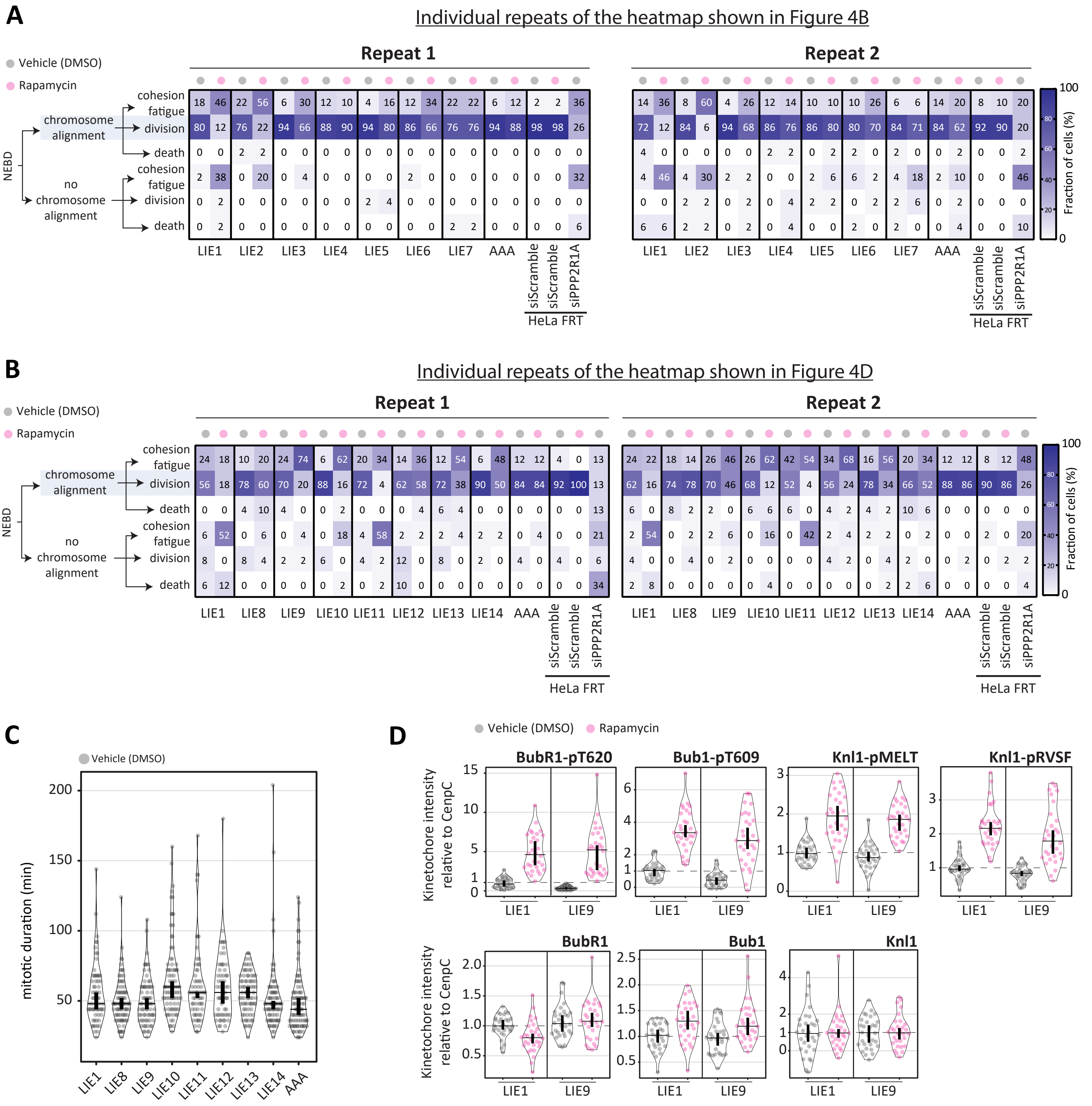


**Supplementary Figure 3 (related to Figure 4) A-B.** The two heatmaps show the frequencies of the cell fates shown in the 2 experimental repeats from the data in Figure 4B (A) and Figure 4D (B). **C.** Mitotic duration – measured as the time between NEBD and chromosome segregation – of the vehicle-treated cells dividing after nuclear envelope breakdown (NEBD) shown in 4D. 59-93 cells from 2 experiments. **D.** Levels of BubR1-pT620, Bub1-pT609, Knl1-pMELT, Knl1-pRVSF, BubR1, Bub1 and Knl1 at unattached kinetochores, in nocodazole-arrested HeLa FRT cells expressing directSLiMs$^{LIE1/LIE9}$ and treated with vehicle or rapamycin for 20min. Note that the distributions of the vehicle-treated directSLiM$^{LIE1}$ and directSLiM$^{LIE9}$ cells are also shown in Figure 4E. Kinetochore intensities from 30 cells, 3 experiments.

Data information: Kinetochore intensities in D are normalized to LIE1 vehicle condition. Violin plots show the distributions of mitotic duration (C) or the distributions of kinetochore intensities between cells (D). For each violin plot, each dot represents an individual cell, the horizontal line represents the median and the vertical one the 95% CI of the median, which can be used for statistical comparison of different conditions (see Materials and Methods).

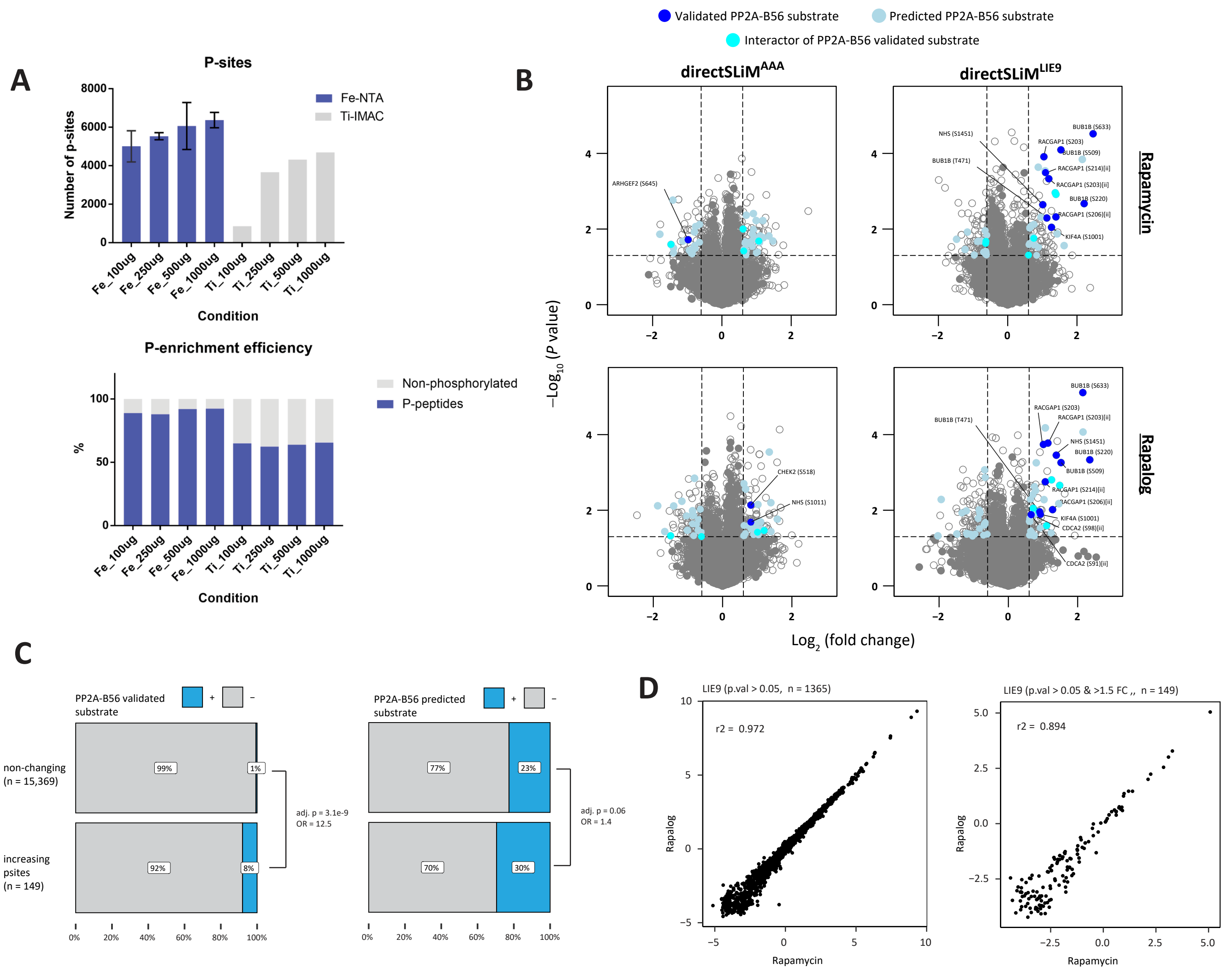


**Supplementary Figure 4 (related to Figure 5). A.** Comparison of Ti-IMAC beads versus Fe-NTA columns for phosphopeptide enrichment. Different amounts of phosphopeptides from mitotic HeLa-FRT cells were subjected to enrichment by these two methods. Fe-NTA showed the highest number of detected phosphorylation sites with all different loading amounts (top). Fe-NTA also displayed a higher selectivity for phosphopeptides when compared to Ti-IMAC beads (bottom). **B.** Volcano plot showing upregulated phosphorylation sites after treatment with rapamycin or rapalog in directSLiMs$^{LIE9}$ cells. Solid dots are either validated substrates, predicted substrates or interactors of validated PP2A-B56 substrates (see methods for details). Only hits above a -Log10(P value = 0.05) and >1.5-fold change are color coded. **C.** Assessing enrichment of PP2A-B56 validated (left) and predicted (right) substrates in proteins with increasing phosphorylation sites. D. Scatter plot showing correlation of rapamycin and rapalog treatment in all changing sites (left) and in the top changing sites (right) in directSLiMLIE9 cell line.

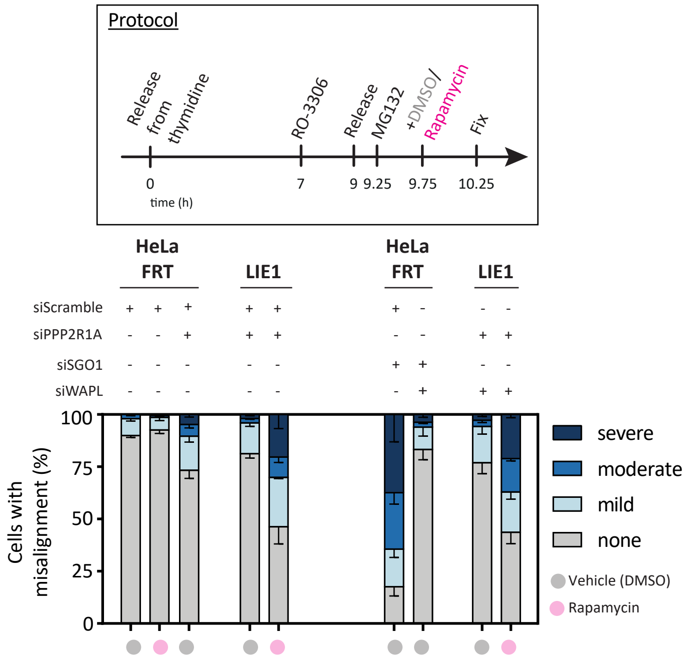


**Supplementary Figure 5 (related to Figure 6).** Evaluating the effects on chromosome alignment in HeLa FRT cells with/without directSLiM[LIE1] expression and treated with vehicle of rapamycin. Top panel: protocol used to visualise chromosome alignment in fixed samples (see Materials and Methods for details). Bottom panel: graph showing mean frequencies of chromosome misalignment (±SEM) of 3 experiments, 100 cell quantified per condition per experiment.
